# Mesenchymal-derived neural progenitors underlie local *insulin* production and neuronal transdifferentiation during retina regeneration

**DOI:** 10.64898/2026.05.20.726251

**Authors:** Bidhi Diwedi, Anindita Neog, Aissette Baanannou, Pritha Das, Romain Menard, Caroline Halluin, Dexter Morse, Kevin Emmerich, James H. Thierer, Michael Patnaude, Frederic Bonnet, Joel H. Graber, Jeff S. Mumm, Romain Madelaine

## Abstract

In humans, retinal-neuron death, optic-nerve injuries, and associated neurodegenerative diseases, such as glaucoma and age-related macular degeneration, often lead to permanent vision loss. While the capacity for regeneration is low in the human nervous system, including the retina, some non-mammalian vertebrate species, including zebrafish, are capable of endogenous neuronal regeneration after injury. Unlike mammals, zebrafish do not form a scar that inhibits axonal and neuronal regeneration after injury. Rather, they harbor neural progenitor and stem-cell populations allowing regeneration of entire parts of the nervous system and restoration of tissue integrity and function. In the zebrafish retina, cycling neural progenitor cells of the ciliary marginal zone and quiescent resident neural stem cells (the latter of which are also called Müller glial cells) participate in neuronal regeneration following different types of injury. In this study, we report the identification of a novel, additional cellular source participating in neuronal regeneration of neurons in the zebrafish retina after genetic ablation of retinal ganglion cells. Before injury, these progenitor cells express molecular markers of neural-crest-cell and/or fibroblast identity, such as *sox10*, *pdgfrb*, and *eya2*, while after neuronal ablation they also express proneural factors including the *ascl1a* and *olig2* genes. Combining genetic ablation of neurons with photoconversion or Cre/Lox-dependent genetic lineage tracing of *sox10*-expressing cells, we demonstrated that these cells can differentiate into post-mitotic retinal neurons in the ganglion cell layer (GCL) in the absence of cell proliferation. We also showed, surprisingly, that this progenitor population locally produces *insulin* mRNA, and that insulin signaling is involved in the accumulation of mesenchymal-derived neural progenitors in the GCL and in their subsequent transdifferentiation into RGCs. This work reveals an unexpected and novel cellular mechanism of transdifferentiation, dependent on a neural-crest-derived mesenchymal cell population, participating in neuronal regeneration in the zebrafish retina. The discovery of this plastic cell population could potentially lead to new strategies to promote the formation of new neurons in the mammalian retina.

## INTRODUCTION

In humans, damage to retinal neurons and the optic nerve, as observed in conditions like glaucoma and age-related macular degeneration, often results in irreversible vision loss. These diseases primarily affect retinal ganglion cells (RGCs) and photoreceptors, respectively ^1–4^ leading to progressive neurodegeneration and permanent loss of function. Currently, there are no effective clinical strategies to restore vision once retinal neurons are lost. A major challenge in regenerative medicine is to identify safe and effective methods to replace these neurons and re-establish functional retinal circuits.

Stem cell-based approaches offer a promising path toward neural repair, as they can self-renew and differentiate into multiple lineages^5,6^. Transplantation of exogenous stem cell-derived neurons has shown the potential to integrate into host tissue^7–10^. However, this approach faces significant challenges and risks^11–13^, including immune rejection, limited integration in the neuronal network, difficulty in directing precise cell fate after transplantation, and vision loss. Several recent studies have shown that endogenous neural stem cells (NSCs) can differentiate under certain conditions into neurons and glia, and have been explored as potential sources of neuron replacement in the retina^14–20^. In the future, the use of endogenous progenitors for cell reprogramming may open new avenues for tissue regeneration.

While regenerative capacity is limited in the mammalian central nervous system^6,21^, several non-mammalian vertebrates, notably zebrafish, possess robust neural regenerative abilities^22–24^. Unlike mammals, zebrafish do not form fibrotic and glial scars that inhibit tissue regeneration^25,26^. Instead, in some animals such has the zebrafish, they retain a population of resident retinal stem cells, i.e., Müller glial cells (MGs) in the retina, that support regeneration and restoration of function, i.e., vision in the retina, following injury^21–24^. Under physiological conditions, MGs remain quiescent to prevent stem-cell-pool depletion. Upon injury, these cells can re-enter the cell cycle and regenerate various types of retinal neurons. In zebrafish, this reactivation involves re-expression of proneural transcription factors such as *ascl1a sox2*, which are both necessary and sufficient for MG-dependent regeneration of retinal neurons^27–30^

Although MGs in mammals typically respond to injury with reactive gliosis, they exhibit latent neurogenic potential. Under certain conditions *in vitro*, mammalian MGs can differentiate into photoreceptors and bipolar cells^31–33^. Notably, forced expression of *Ascl1* in murine MGs can promote neurogenesis following injury, generating amacrine, bipolar, and photoreceptor neurons. However, this capacity has been largely limited to young animals, due to restricted chromatin accessibility in adult MGs; yet, epigenetic modulation, such as histone deacetylase inhibition, can partially overcome these barriers and enhance regeneration^18,19^. Recently, studies have shown that the combination of multiple transcription factors (e.g., *Ascl1*, *Atoh1*, *Pou4f2*) has potential in specifying a broader range of retinal neuron types, including RGC-like cells^14,20,34^. Nonetheless, the retinal-neuron regenerative capacity remains insufficient to treat retinal degeneration diseases and restore vision in patients, highlighting the need to identify alternative or complementary molecular and/or cellular regenerative mechanisms.

To investigate neuronal regeneration in a naturally regenerative system, we employed zebrafish models using the nitroreductase/metronidazole system for targeted genetic ablation of retinal neurons^35–37^. Surprisingly, after ablation of RGCs, we identified a regenerative response that appears independent of that of MGs. Specifically, we discovered a previously uncharacterized population of mesenchymal-derived progenitor cells with neural-crest-like features contributing to RGC regeneration. Through transcriptomic analysis, genetic lineage tracing, and functional assays, we determined that this population, marked by the expression of genes including *sox10*, *pdgfrb*, *acta2*, and *eya2*, exhibits a neural progenitor identity. Indeed, after RGC ablation, these cells accumulate in the ganglion cell layer (GCL) and express proneural markers such as *ascl1a* and *olig2*, indicating acquisition of a proneural cell identity. Lineage-tracing analysis using photoconversion and Cre/LoxP dependent recombination revealed that *sox10*/*pdgfrb+* cells give rise to RGCs. Interestingly, this regenerative process does not require cell-cycle activation and proliferation, suggesting that these cells undergo transdifferentiation and a direct conversion into neurons. In addition, we demonstrated that communication pathways involved in fibroblast activation such as Pdgf, Wnt, Tgf-β, Bmp, and Hippo signaling are involved in this process. Finally, inhibition of Myosin II activity and photoconversion analysis suggested that these cells migrate from an anterior mesenchyme structure of the eye into the GCL, supporting this newly identified process of neuronal regeneration.

Unexpectedly, we determined that *insulin* mRNA is expressed in *pdgfrb+* neural progenitors during RGC regeneration. Although *insulin* is classically known to be produced in the pancreas^38,39^, growing evidence from model organisms suggests extra-pancreatic expression in several tissues^40,41^, including the retina^42^. In our regenerative zebrafish model, pancreatic-derived insulin is dispensable while global inhibition of insulin signaling impairs RGC regeneration, suggesting an important role for the local production of *insulin* mRNA by mesenchymal-derived progenitors in the neuro-regenerative response.

In conclusion, this work provides compelling evidence that a cranial neural-crest-derived population of progenitor cells contributes to RGC regeneration. Previous studies of regenerative species such as zebrafish have demonstrated that neural-crest- and mesenchymal-derived cells have the differentiation and re-differentiation potential to repair tissues including the heart, kidney, and limb^43–50^, Our finding identifies that mesenchymal-derived cells can regenerate retinal neurons through transdifferentiation, a process modulated by local production of *insulin* mRNA, a discovery that opens a new avenue for regenerative therapy. In mammals, where MGs may have a limited regenerative ability, targeting similar plastic mesenchymal or fibroblast cell populations may offer a viable strategy to restore vision.

## RESULTS

### *sox10-*expressing cells accumulate in the retina after genetic ablation of RGCs

To identify new genes, and potentially new cellular mechanisms, involved in retinal neuron regeneration after injury, we used a previously published single-cell RNAseq dataset established after genetic ablation of retinal ganglion-cells (RGCs) in zebrafish^36^. We refined the analysis on a subset of cell populations by focusing on specific neural and neuronal cells, but also including specific glial and vascular markers, as these populations have not been scrutinized in the study mentioned above (Fig. 1A and Fig. S1A). As expected, neural progenitors, RGC precursor cells, and post-mitotic neurons are the cell types highly represented in the subset analysis, while vascular cells and myelinated oligodendrocytes are rarely identified and not identified, respectively. We next inferred a common analysis for control and ablated samples (24 hours after ablation) at 6 days post fertilization (dpf). We observed heterogeneity in cell identity as indicated by splitting of the neural progenitor cells into multiple cell clusters, possibly reflecting the expression of different proneural factors (Fig. 1A and Fig. S1A). Interestingly, we identified small populations of cells expressing the *sox10* gene, potentially of cranial neural-crest or glial-cell identity, in two different clusters, presenting evidence of cell-transition events (Fig. 1A and Fig. S1A, B). Cells expressing *sox10* are subclustered at the arm of the main cluster, for both cluster 1 and cluster 2. Pseudotime trajectory and cell-velocity analyses suggest that *sox10*+ cells in clusters 1 and 2 could participate in the establishment of a neural progenitor population after RGC ablation (Fig. 1A and Fig. S1A, B). Based on this observation, we sought to investigate the potential role of *sox10*-expressing cell-derived progeny during neuronal regeneration in the zebrafish retina.

**Figure 1.**
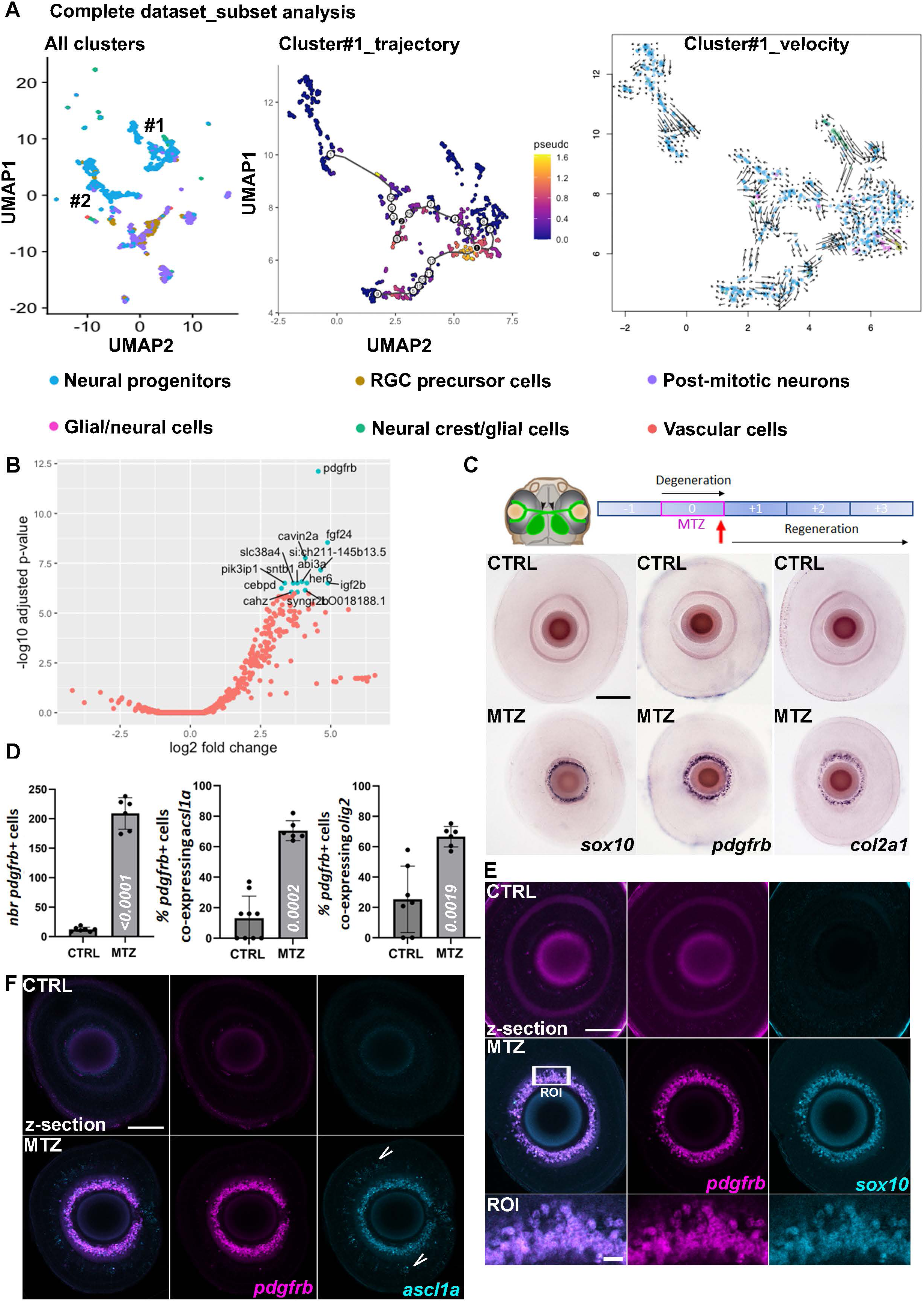
*sox10*-expressing cells are neural progenitors accumulating in the GCL after RGCs genetic ablation. **(A)** Sub-analysis of single-cell RNAseq dataset at 6 dpf (days post-fertilization) after genetic ablation of RGCs using Tg(*islet2b:gal4*); Tg(*uas:NTR-yfp*) with MTZ treatment between 5 and 6 dpf. UMAPs show the clustering of various retinal cell types. Each cluster is color-coded to represent different cells as indicated in the figure. The left panel underscores the fate proximity of *sox10*+ cells (green) with neural progenitor cells (blue) in two clusters. Trajectory (middle panel) and cell velocity (right panel) in cluster 1 reveal a pattern of movement from *sox10*-expressing cells to young and mature neurons. **(B)** Differential gene-expression analysis of *sox10*+ cells against all other cells in the cluster 1, indicating genes significantly associated with *sox10* expression in cluster 1. **(C)** Analysis of *sox10*, *pdgfrb*, and *col2a1* mRNA expression by *in situ* hybridization in control or MTZ-treated (between 5 and 6 dpf) larvae (Tg(*islet2b:gal4*); Tg(*uas:NTR-yfp*)) at 6 dpf. **(D)** Quantification of the number of *pdgfrb*+ cells in the GCL at 6 dpf after genetic ablation of RGCs between 5 and 6 dpf (control; n=7 and MTZ; n=6; left). Analysis of the percentage of *pdgfrb*+ cells expressing the proneural factors *ascl1a* (control; n=9 and MTZ; n=6; middle) and *olig2* (control; n=7 and MTZ; n=6; right). Statistical significance is determined by Mann-Whitney test, non-parametric. **(E)** Representative image (confocal z-section) of *sox10* and *pdgfrb* co-expression in the GCL after RGC genetic ablation (between 5 and 6 dpf) using HCR fluorescent *in situ hybridization*. **(F)** Representative image (confocal z-section) of *pdgfrb* and *ascl1a* co-expression in the GCL after RGC genetic ablation (between 5 and 6 dpf) using HCR fluorescent *in situ hybridization*. Arrows indicate inner *ascl1a* expression outside of the GCL. Error bars represent s.d. Scale bars: 100 μm.

To study retinal-neuron regeneration in zebrafish larvae and the potential role of the *sox10*+ cell populations during this process, we took advantage of an innovative and previously established combination of transgenic lines^36^ to perform specific genetic ablation of RGCs: Tg(*islet2b*:gal4); Tg(*uas*:NTR2.0-yfp). Following the addition of a drug (metronidazole; MTZ) in the medium (24 hours at 10 μM), neurons expressing nitroreductase (NTR2.0) will die by apoptosis^35^. Ablation of RGCs was performed between 5 and 6 dpf (Fig. S1D). We observed a potent reduction in expression of genes associated with RGC identity, including *islet2b*, *pou4f1, pou4f2, pou4f3*, and *elavl4*, via RNAseq analysis (Table S1), indicating a significant reduction in the number of cells expressing these genes. We validated the above result with fluorescent imaging and confirmed the previous results^36^ that demonstrated RGC and optic-nerve degeneration after 24 hours of MTZ treatment (Fig. S1D, E). Supporting a degenerative phenotype, we also observed an inflammatory response and accumulation of Tg(*mpeg:mCherry)*+ microglia in the GCL after neuronal ablation (Fig. S1F). We next validated our bioinformatic predictions and revealed *sox10* gene expression in the retina after genetic ablation of RGCs (1-day post-ablation; dpa) by *in situ* hybridization (Fig. 1C). However, the putative function of the *sox10* gene and the role of *sox10*-expressing cells during retinal-neuron regeneration still need to be investigated.

### *sox10*-expressing cells carry a proneural identity and accumulate in the GCL independently of cell proliferation

Because transcriptomics analysis and *in situ* hybridization (ISH) analyses revealed a potential role for *sox10*-expressing cells in retinal-neuron regeneration, we next performed differential expression analysis of *sox10*-expressing cells against all other cells in the cluster 1 or 2 to identify genes significantly enriched and associated with *sox10* expression (Fig. 1B and S1C). Interestingly, clusters 1 and 2 share the same top gene associated with *sox10* expression and neural-crest and/or glial-cell identity: *pdgfrb*. We validated the increased expression of the *pdgfrb* gene, as well as of the *col2a1* gene, which is highly associated with Pdgfrb+ cell identity in different tissues (Fig. 1C). We quantified an accumulation of ∼200 *pdgfrb*+ cells in the GCL at 6 dpf and demonstrated co-expression of the *sox10* and *pdgfrb* genes by hybridization chain reaction (HCR) fluorescent *in situ* (Fig. 1D, E). We determined that *pdgfrb*+ cells accumulate in the GCL but lack expression of the marker of post-mitotic RGCs, HuC/D (Fig. S1G). To better characterize *sox10*/*pdgfrb*-expressing cells under homeostatic and ablated conditions, we analyzed cell proliferation by EdU incorporation in *sox10:kaede*+ cells and revealed that while we observed EdU incorporation in the retina (potentially in neural progenitor cells derived from the ciliary marginal zone and/or MGs), and an accumulation of *sox10*+ cells in the GCL, we did not detect evidence of EdU incorporation in *sox10*+ cells (Fig. S1H). Together, these experiments indicate that cell proliferation is induced after ablation of RGCs, but that the proliferative response is likely not responsible for the accumulation of *sox10*/*pdgfrb*-expressing cells in the GCL.

To investigate whether *sox10/pdgfrb*+ cells are undergoing a transition to a neural-progenitor identity, we analyzed the expression of some genes that encode proneural factors known to control neurogenesis and neuronal differentiation^51^. To characterize a potential cellular process of neuronal differentiation involving *pdgfrb*-expressing cells and proneural gene expression, we performed HCR *in situ* hybridization at 6 dpf and showed that the *ascl1a* and *olig2* genes, but not the *neurod1* gene, are robustly expressed 1 day after RGC ablation (Fig. 1F and Fig. S2B, C). Interestingly, after RGC ablation, *ascl1a* expression is detected not only in the GCL, but also in medial and distal layers of the retina, suggesting that cells that are not expressing *pdgfrb*/*sox10* (potentially neural progenitor cells derived from the ciliary marginal zone and/or MGs) are also involved in expansion of *ascl1a* gene expression. We calculated that ∼70% of *pdgfrb*+ cells also express *ascl1a*, ∼65% express *olig2*, and only *∼*1% co-express *neurod1* (Fig. 1D and Fig. B’), indicating that, in this context of RGC degeneration, a significant number of *pdgfrb+* cells also carry a neural progenitor identity. Together, these data indicate that a neuronal regenerative response may arise from cells expressing the *sox10* and *pdgfrb* genes after genetic ablation of RGC in the zebrafish retina.

### *sox10*-expressing progenitor cells transdifferentiate into retinal neurons

Our data suggest that in response to RGC ablation, a cell population in the zebrafish retina expresses *sox10*, *pdgfrb*, and some proneural genes, and accumulates in the retinal GCL. Intriguingly, it appears that modification of the gene-expression profile in these cells occurs independently of cell-proliferation activation, suggesting that retinal progenitors of the ciliary marginal zone, or derived from MGs, are not the major participants in the establishment of neural progenitors of a potential neural-crest origin. Instead, we propose that an additional cellular mechanism of transdifferentiation occurs after RGC ablation in the zebrafish retina. Cellular plasticity may be achieved in neural-crest-derived progenitors under the activity of transcription factors such as Sox10, Olig2, and Ascl1a.

To determine whether *sox10*/*pdgfrb*+ cells can differentiate into retinal neurons, we performed photoconversion and genetic cell lineage tracing of *sox10-* and/or *pdgfrb*-expressing cells. We used Tg(*sox10:Kaede*) and Tg(*pdgfrb:dendra2*) to perform photoconversion of *sox10-*and *pdgfrb*-expressing cells, respectively (photoconversion from green to red fluorescence using UV light; Fig. S2C). When we performed photoconversion before genetic ablation of neurons (between 4 and 5 dpf), we determined that ∼1% of PhotoConverted (PC)-Kaede+ cells in the GCL incorporated EdU at 6 dpf (Fig. 2A, A’). This result supports our previous observations that cell proliferation does not play a major role in the accumulation of *sox10*/*pdgfrb*+ cells in the GCL. In addition, we demonstrated that after RGC ablation, ∼98% of PC-Kaede+ or PC-Dendra2+ cells in the GCL are neurons expressing HuC/D at 9 dpf (Fig. 2B-B”). These observations suggest that a *sox10*+ cell population present in the zebrafish retina has the ability to transdifferentiate into neurons. To validate the above hypothesis, we performed concomitant photoconversion of Kaede or Dendra2 proteins with the ablation of RGCs between 5 and 6 dpf, followed by EdU incorporation between 6 and 7 dpf. We demonstrated that the majority of HuC/D+ neurons derived from *sox10:kaede*+ cells have not incorporated EdU at 9 dpf (Fig. 2C, C’). Finally, genetic cell lineage tracing using *sox10*:CreERT2-dependent LoxP-site recombination confirms our previous observation that *sox10*-derived cells are involved in regeneration of RGCs in the zebrafish retina (Fig. 2D). Altogether, these data support the concept of a *sox10*+ cell population responding to genetic ablation of RGCs by accumulating in the injured GCL to acquire a proneural identity and differentiate into neurons.

**Figure 2.**
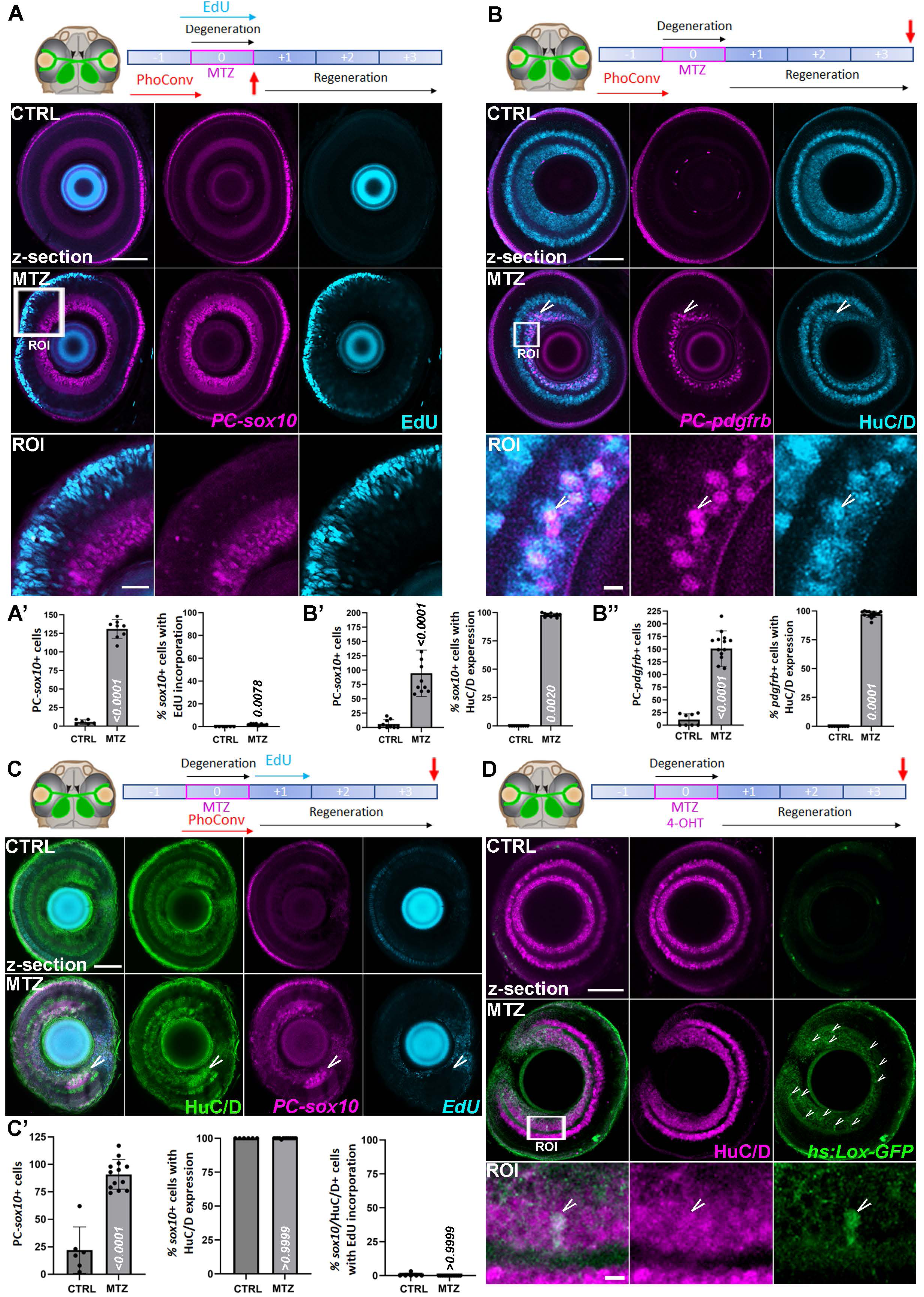
***sox10*-expressing cells transdifferentiate into RCCs without cell proliferation. (A)** Representative image (confocal z-section) of cell lineage tracing analysis of *sox10*+ cell fate at 6 dpf by Kaede photoconversion (PC using Tg(*sox10:Kaede*) between 4 and 5 dpf) followed by genetic ablation of RGCs using (Tg(*islet2b:gal4*); Tg(*uas:NTR-yfp*)) with concomitant EdU incorporation (between 5 and 6 dpf). **(B)** Representative image (confocal z-section) of cell lineage tracing analysis of *pdgfrb*+ cell fate at 9 dpf by Dendra2 photoconversion (PC using Tg(*pdgfrb:dendra2*) between 4 and 5 dpf) followed by genetic ablation of RGCs using Tg(*islet2b:gal4*); Tg(*uas:NTR-yfp*) between 5 and 6 dpf. White arrow indicates photoconverted cells expressing HuC/D, a marker of differentiated RGCs. **(A’)** Quantification of the number of PC-*sox10*+ cells in the GCL at 6 dpf after genetic ablation of RGCs between 5 and 6 dpf (control; n=6 and MTZ; n=8; Mann-Whitney test, non-parametric; left). Analysis of the percentage of PC-*sox10*+ cells incorporating EdU (control; n=6 and MTZ; n=8; one-sample Wilcoxon test; right). **(B’)** Quantification of the number of PC-*sox10*+ cells in the GCL at 9 dpf after genetic ablation of RGCs between 5 and 6 dpf (control; n=9 and MTZ; n=10; Mann-Whitney test, non-parametric; left). Analysis of the percentage of PC-*sox10*+ cells expressing HuC/D (control; n=9 and MTZ; n=10; one-sample Wilcoxon test; right). **(B’’)** Quantification of the number of PC-*pdgfrb*+ cells in the GCL at 9 dpf after genetic ablation of RGCs between 5 and 6 dpf (control; n=8 and MTZ; n=14; Mann-Whitney test, non-parametric; left). Analysis of the percentage of PC-*pdgfrb*+ cells expressing HuC/D (control; n=8 and MTZ; n=14; one-sample Wilcoxon test; right). **(C)** Representative image (confocal z-section) of cell lineage tracing analysis of *sox10*+ cell fate at 9 dpf by Kaede photoconversion using Tg(*sox10:Kaede*) with concomitant genetic ablation of RGCs (between 5 and 6 dpf) followed by EdU incorporation (between 6 and 7 dpf). White arrow indicates photoconverted cells expressing HuC/D, a marker of differentiated RGCs. **(C’)** Quantification of the number of PC-*sox10*+ cells in the GCL at 9 dpf after genetic ablation of RGCs between 5 and 6 dpf (control; n=6 and MTZ; n=13; t-test, parametric, two-tailed; left). Analysis of the percentage of PC-*sox10*+ cells (control; n=6 and MTZ; n=13; one-sample Wilcoxon test) expressing HuC/D (middle) and/or incorporating EdU (right). **(D)** Representative image (confocal z-section) of cell lineage tracing analysis of *sox10*+ cell fate at 9 dpf by creERT2-dependent loxP-site recombination (using 4-OHT with Tg(*sox10:creERT2*); Tg(*hs*:*loxp-rfp-loxp-gfp*)) with concomitant genetic ablation of RGCs (using Tg(*islet2b:gal4*); Tg(*uas:NTR-mCherry*) between 5 and 6 dpf). White arrows indicate *sox10*-derived cells expressing HuC/D, a marker of differentiated RGCs. Error bars represent s.d. Scale bars: 100 μm or 50 μm in ROI.

### *sox10*-expressing neural progenitors also carry a mesenchymal identity

We next tested whether *sox10* gene function is critical for establishment of the *sox10*/*pdgfrb*-expressing neural progenitor population in the GCL. Using HCR *in situ*, we analyzed *pdgfrb* expression in the *sox10* mutant. We observed that accumulation of *pdgfrb*+ cells was not significantly affected in the *sox10* loss-of-function genetic background (Fig. S2D, D’), indicating that the activity of this gene is dispensable for the accumulation of *pdgfrb+* progenitor cells after RGC ablation in the zebrafish retina. Similarly, loss of *pdgfrb* function does not affect establishment of *sox10*+ cells in the GCL after genetic ablation of RGC (Fig. S2D, D’). We reasoned that *pdgfra* could have a redundant function with *pdgfrb* in the *sox10*+ cell population. To test this hypothesis, we analyzed *pdgfra* expression and determined that, similarly to the *pdgfrb* gene, its mRNA is detected in the GCL after genetic ablation of neurons (Fig. 3A). We next used a well-characterized PDGFR-inhibitor^52^ and demonstrated that Pdgfr signaling is required for the accumulation of *sox10*+ cells in the GCL after RGC ablation (Fig. 3B, B’).

**Figure 3.**
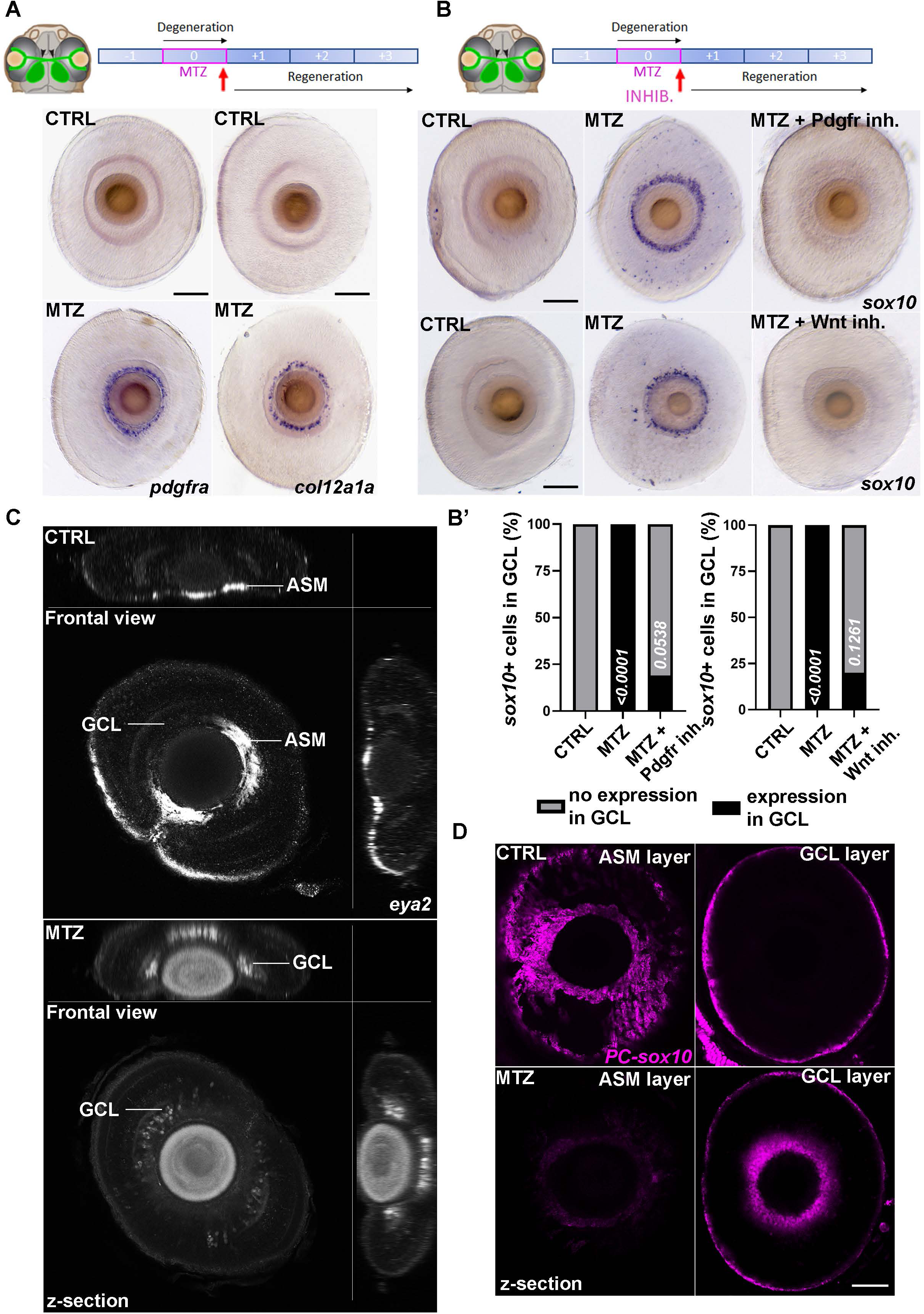
*sox10*+ cells are fibroblast-like cells potentially derived from the anterior segment mesenchyme. **(A)** Analysis of *pdgfra* and *col12a1a* mRNA expression by *in situ* hybridization in control or MTZ-treated (between 5 and 6 dpf) larvae (Tg(*islet2b:gal4*); Tg(*uas:NTR-mCherry*)) at 6 dpf. **(B)** Analysis of *sox10* mRNA expression by *in situ* hybridization in control or MTZ-treated (between 5 and 6 dpf) larvae (Tg(*islet2b:gal4*); Tg(*uas:NTR-mCherry*)) at 6 dpf with or without inhibition of the Pdgfrb or Wnt signaling pathway. **(B’)** Quantification of the number of retina harboring *sox10*+ cells in the GCL at 6 dpf after genetic ablation of RGCs between 5 and 6 dpf using Fisher’s exact test (control; n=22 MTZ; n=11; MTZ + Pdgfrb inhibitor; n=26; left, and control; n=14 MTZ; n=18; MTZ + Wnt inhibitor; n=20; right). **(C)** Representative image (confocal z-section) of *eya2* expression in the ASM or GCL layer after RGC genetic ablation (between 5 and 6 dpf) using HCR fluorescent *in situ hybridization*. Without genetic ablation of RGCs, *eya2* expression is detected in the front of the eye, above the GCL layer. After ablation of RGCs, *eya2* expression is also detected below the front of the eye, likely in the injured GCL. **(D)** Representative image (confocal z-section) of *PC-sox10:kaede* expression in the ASM or GCL layer after RGC genetic ablation (between 5 and 6 dpf). Without genetic ablation of RGCs, *PC-kaede* expression is detected in the front of the eye, above the GCL layer. After ablation of RGCs, expression is reduced in front of the eye but detected below it, likely in the injured GCL. Scale bars: 100 μm.

The above result as well as the result for *col2a1* gene expression after ablation (Fig. 1C), raised the hypothesis that *sox10*/*pdgfrb*+ cells are neural-crest-derived fibroblasts that are activated after injury. To test this possibility, we analyzed the expression of other genes associated with fibroblast-like identity during regeneration. We observed that *col12a1*, a well-characterized pro-regenerative collagen gene^52,53^, as well as *acta2,* a marker of activated fibroblasts, are expressed in the GCL after RGC ablation (Fig. 3A and Fig. S2E). This observation supports the hypothesis that *sox10*/*pdgfrb*+ cells accumulating in the GCL after injury are fibroblast-like cells from a neural-crest origin. We next investigated whether this previously uncharacterized process of neuronal regeneration is independent of the classical resident retinal stem cell population, i.e., MGs. We performed concomitant ablation of RGCs and MGs using the Tg(*islet2b:gal4*); Tg(*uas:NTR2.0-mCherry*); Tg(*gfap:NTR2.0-tagRFP*) lines and demonstrated that genetic ablation of *gfap*+ cells has no impact on accumulation of *sox10*+ progenitors in the GCL after RGC degeneration (Fig. S3A). This result supports our previous observations that this process of progenitor accumulation in the GCL is independent of cell proliferation and relies on a newly identified cellular source of neural progenitors.

### The anterior segment mesenchyme is the source of *sox10*-derived neural progenitors

The data above led us to hypothesize that a mesenchymal/fibroblastic cellular source underlies a new mode of neuronal regenerative response in the zebrafish retina. Supporting this hypothesis, HCR *in situ* revealed co-expression of the *eya2* gene with *pdgfrb*+ cells in the GCL (Fig. S3B), indicating that *sox10*+ neural progenitors also carry a mesenchymal identity. The co-expression of *pdgfrb* and *eya2* prompted us to propose that the anterior segment mesenchyme (ASM) of the zebrafish retina is, at least partially, the cellular source of these newly identified progenitors. Indeed, this structure is known to derive from Sox10+ cranial neural crest cells during embryonic development, and some of the marker genes tested above, e.g., *sox10* and *eya2*, have been associated with development of the ASM, which will later form some of the anterior structures of the eye such as the stroma of the cornea, of the iris, and of the ciliary body^54–57^. We observed that after RGC ablation, the anterior domain of *eya2* expression surrounding the lens is reduced, but it expands in the GCL located below (Fig. 3C). We hypothesized that retinal mesenchymal cells from the ASM are activated and accumulate in the GCL after injury, as suggested by the dynamic pattern of *eya*2 expression. To confirm the above hypothesis, we used the *sox10:kaede* line to perform photoconversion between 4 and 5 dpf (Fig. S2C), followed by RGC ablation between 5 and 6 dpf. We observed that following targeted ablation of neurons, *PC-Sox10* expression is strongly detected in the GCL (CTRL: n=0/15; MTZ: n=12/12 with expression in the RGC layer; Fig. 3D). Because, after genetic ablation of RGCs, *PC-Sox10* expression is detected in the GCL and reduced in the ASM layer surrounding the lens, we proposed that *sox10*+ cell accumulation in the GCL is dependent primarily on plastic cells that are derived from the retinal ASM and that express the *eya2* gene. Altogether, these data support the concept that a cellular source of mesenchymal origin participates in a newly identified mode of neuronal regeneration in the zebrafish retina.

### Signaling pathways regulating fibroblast activation are involved in the regenerative process dependent on *sox10*+ progenitors

Because pharmacological inhibition of Pdgf signaling impaired the accumulation of *sox10+* neural progenitors in the GCL, we next asked whether other communication pathways involved in fibroblast activation are required to support accumulation of *sox10*+ cells in the GCL after injury. Similarly to Pdgf inhibition, blocking Wnt, TGF-β, or BMP signaling limits the accumulation of *sox10*+ cells in the GCL (Fig. 3B, B’, and S3C), suggesting that these pathways are involved in the neuronal-regeneration process dependent on mesenchymal-derived neural progenitors. In addition, inhibition of myosin II indicates that cellular motor activity is necessary for the accumulation of *sox10*+ progenitors in the GCL after RGC ablation (Fig. S3C). Finally, we used a selective inhibitor of glycogen synthase kinase 3 (GSK-3) to activate the canonical Wnt signaling pathway with concomitant ablation of RGCs, and we identified that it leads to the detection of ectopic *pdgfrb*+ cell outside of the GCL, supporting our previous observation that Wnt signaling is involved in the regenerative response dependent on mesenchymal-derived cells (Fig. S3D). Because our analysis suggests that cell proliferation is not the primary mechanism involved in this regenerative process, we propose that cell motility and migration from the ASM to the site of injury in the GCL regulates the neuronal-regeneration process associated with mesenchymal-derived neural progenitors. Together, these results suggest that reducing cellular contractile capacity affects the accumulation of mesenchymal-derived progenitors in the GCL and impairs regenerative neurogenesis.

### Local *insulin* production by mesenchymal-derived neural progenitors impacts regeneration

We performed GO and KEGG analyses of the RNAseq data comparing control and ablation conditions, and identified that neurogenesis/neurodifferentiation processes and insulin signaling are upregulated following genetic ablation of RGCs (Fig. S4B, B’). This observation suggested a potential role for insulin signaling in the regenerative process. While insulin is synthesized predominantly in the pancreas, emerging evidence suggests that *insulin* mRNA is also expressed in extra-pancreatic tissues^40,41^. In the mammalian retina, insulin mRNA expression was recently detected in retinal pigment epithelium (RPE), with a role in phagocytosis of damaged photoreceptors^42^. In the chicken or zebrafish retina, exogenous application of insulin stimulates cell proliferation during regeneration after injury^58,59^. Strikingly, we discovered that *insulin* mRNA is expressed in *pdgfrb*-expressing cells following RGC ablation (Fig. S4A) and that these cells also co-express *insulin* and the *ascl1a* proneural gene (Fig. 4A). This finding suggests that local *insulin* production by mesenchymal-derived neural progenitors could be involved in retinal-neuron regeneration. To test this hypothesis, we first performed concomitant ablation of RGCs and *insulin*+ cells using Tg(*ins*:NTR2.0-mCherry; Tg(*islet2b*:NTR2.0-mCherry) lines, and demonstrated that this experimental setup leads to ablation of *insulin*+ beta cells in the pancreas, but not *insulin*+ neural progenitors in the retina (Fig. 4B and Fig. S4C, C’). This experiment also revealed that *insulin* expression from the pancreas is dispensable for *insulin*+ cell accumulation in the GCL after injury. Next, we targeted insulin-receptor activity by using a selective insulin-receptor tyrosine kinase inhibitor^60–62^. This strategy indicated that global loss of insulin-receptor activity affects accumulation of *pdgfra*+ progenitor cells in the GCL (Fig. 4C, C’), potentially impacting the regenerative capacity of the zebrafish retina. We also demonstrated by inhibition of YAP-TEAD interaction that the Hippo pathway, which is also an important component of the metabolic response that can interact with and regulate insulin signaling, is involved in the regenerative process that is dependent on mesenchymal-derived progenitors (Fig. 4C). We next determined that insulin-receptor genes, e.g., *insra* and *insrb*, are expressed widely in the retina, both in the microenvironment and in *pdgfrb*+ progenitor cells (Fig. S4D), suggesting that locally produced insulin can act in both a paracrine and an autocrine manner, while beta cells can provide endocrine-dependent insulin signaling in the injured retina. Interestingly, we did not observe any upregulation of the *insra* and *insrb* genes after RGC ablation (Fig. S4D), suggesting that modulation of insulin signaling after injury in the retina relies primarily on local increased expression of the *insulin* gene. Finally, using an antibody targeting the insulin peptide in zebrafish, we revealed that the local production of *insulin* mRNA leads to the expression of the hormone in the GCL after injury (Fig. 4D), supporting a local role for insulin during retinal neurons regeneration. Altogether, the above results suggest that local *insulin* production in the retina is involved in the process of neuronal regeneration dependent on mesenchymal-derived neural progenitors.

**Figure 4.**
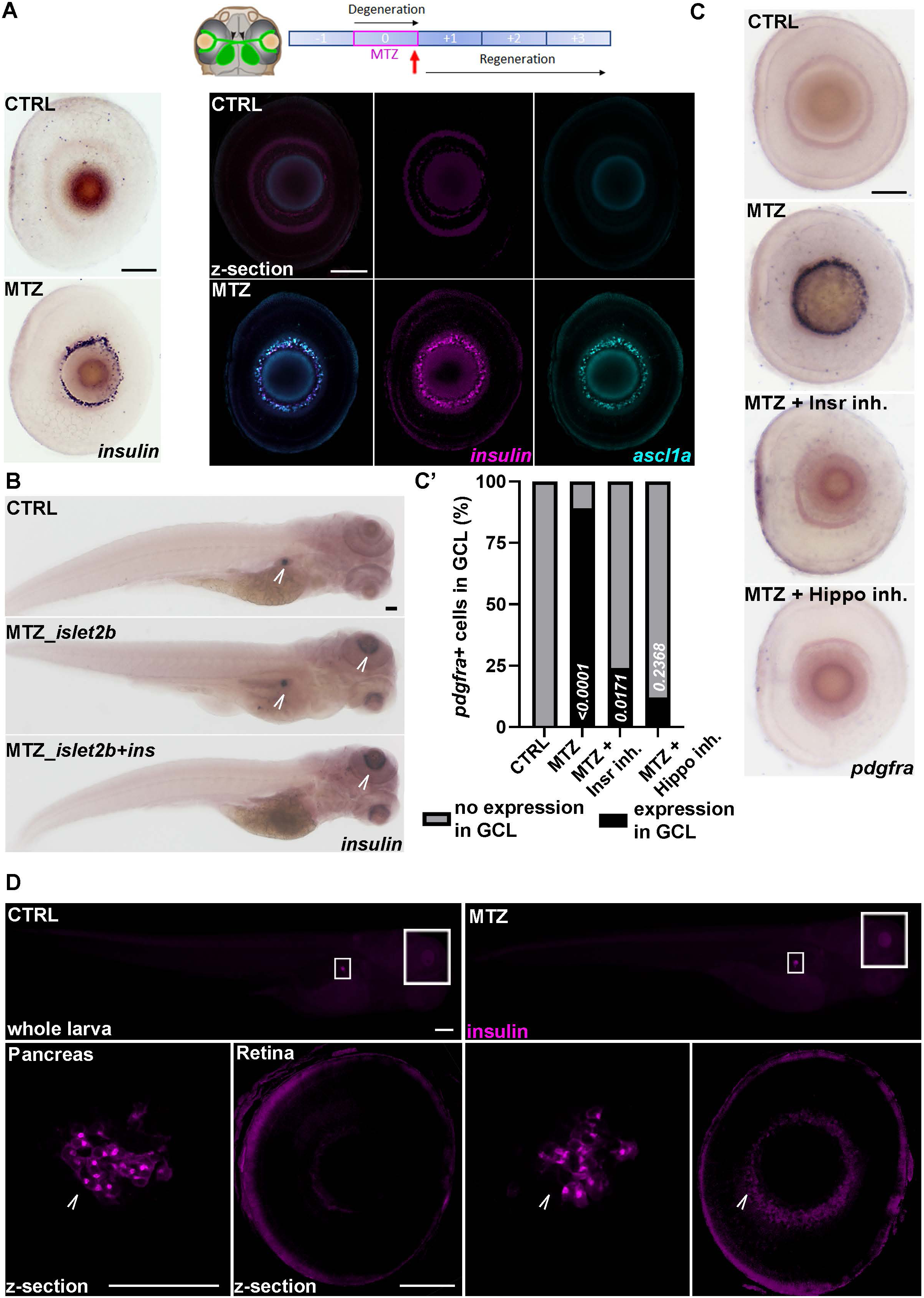
*insulin* mRNA is locally expressed in the retina after ablation of RGCs and Insulin signaling participates in the accumulation of mesenchymal-derived progenitors. **(A)** Analysis of *insulin* mRNA expression by *in situ* hybridization in control or MTZ-treated (between 5 and 6 dpf) larvae (Tg(*islet2b:gal4*); Tg(*uas:NTR-mCherry*)) at 6 dpf. HCR analysis revealed co-expression of the *insulin* and *ascl1a* genes in the GCL after ablation of RGCs. **(B)** Analysis of *insulin* mRNA expression by *in situ* hybridization in control or MTZ-treated (between 5 and 6 dpf) larvae (Tg(*islet2b:gal4*); Tg(*uas:NTR-mCherry*)) and/or Tg(*ins:NTR-mCherry*) at 6 dpf. Arrows show sites of *insulin* expression in the pancreas and/or the retina. **(C)** Analysis of *pdgfra* mRNA expression by *in situ* hybridization in control or MTZ-treated (between 5 and 6 dpf) larvae (Tg(*islet2b:gal4*); Tg(*uas:NTR-mCherry*)) at 6 dpf with or without inhibition of insulin or the Hippo signaling pathway. **(C’)** Quantification of the number of retina harboring *pdgfra*+ cells in the GCL at 6 dpf after genetic ablation of RGCs between 5 and 6 dpf using Fisher’s exact test (control; n=22 MTZ; n=18; MTZ + Insulin inhibitor; n=34; MTZ + Wnt inhibitor; n=25). **(D)** Analysis of insulin peptide expression by immunostaining in control or MTZ-treated (between 5 and 6 dpf) larvae (Tg(*islet2b:gal4*); Tg(*uas:NTR-mCherry*)) at 6 dpf (top: whole larva widefield; bottom; confocal z-section). Arrows show sites of Insulin expression in the pancreas and/or the retina. Scale bars: 100 μm.

## DISCUSSION

Regeneration of retinal neurons in vertebrates remains a major challenge in regenerative biology and medicine. In mammals, neuronal loss caused by optic-nerve injury or degenerative diseases such as glaucoma and age-related macular degeneration is largely irreversible, reflecting the limited intrinsic regenerative capacity of the human central nervous system. In contrast, zebrafish exhibit a remarkable ability to restore retinal structure and function following injury. This regenerative competence has been primarily attributed to Müller glial cells (MGs), which dedifferentiate, proliferate, and generate multipotent progenitors and neurons^21,23,24,63^. Our study expands this paradigm by identifying an unexpected and mechanistically distinct novel cellular source of regenerated neurons.

Our findings indicate that at least three parallel regenerative routes coexist in the zebrafish retina: two well-characterized progenitor populations that undergo proliferative-dependent regenerative neurogenesis, i.e., a population derived from the ciliary marginal zone, and the MG population, and our newly discovered mechanism of neuronal transdifferentiation, potentially originating from mesenchymal-derived cells of the ASM. Such redundancy may contribute to the rapid and robust retinal repair in teleosts, suggesting that regenerative responses are more heterogeneous than previously appreciated. Future work will better define the respective contribution arising from each cellular source to support the process of neuronal regeneration.

The molecular identity of our newly characterized neural progenitor population challenges traditional lineage boundaries. Following neuronal ablation, these cells co-express markers associated with neural-crest and mesenchymal lineages. When accumulated in the GCL, they also acquire expression of proneural transcription factors. These observations align with emerging evidence in other tissues, such as heart, kidney, and limb^43–50,64^, that fibroblast-like cells, in particular those carrying neural-crest features, are not terminally differentiated endpoints but can undergo lineage conversion in response to injury-associated cues.

The signaling pathways regulating our newly identified regenerative process reveal notable overlaps with both regenerative and fibrotic programs. Our pharmacological-inhibition experiments demonstrate that Pdgf, Wnt, Bmp, Tgf-β, and Hippo signaling are required for efficient progenitor accumulation in the GCL of the injured retina. These pathways are well-established regulators of fibroblast activation and extracellular-matrix remodeling, yet they also play central roles in neural development and stem/progenitor-cell maintenance and differentiation ^43^, as demonstrated by examples in regenerative studies, including heart, kidney, or retina, with Wnt and Hippo signaling functioning as shared players across tissues^48,49,65–69^. The involvement of Pdgf, Wnt, Bmp, Tgf-β, and Hippo signaling in this novel regenerative route suggests that retinal injury induces a transient fibroblastic activation program that, in zebrafish, promotes plasticity and regeneration rather than permanent scar formation.

An additional and unexpected regulatory layer is the role of local insulin signaling. Our detection of *insulin* mRNA within *pdgfrb*+ progenitors, coupled with impaired regeneration upon insulin-receptor inhibition, indicates that insulin acts as a key modulator of neuronal regeneration. Pancreatic insulin appears dispensable, whereas local retinal insulin production correlates with progenitor accumulation and positive neuronal regeneration output, suggesting predominantly paracrine and autocrine mechanisms. Insulin signaling may enhance progenitor survival, metabolic fitness, and transcriptional plasticity, integrating metabolic regulation into the regenerative program.

The cellular mechanism we have uncovered in zebrafish shows notable conceptual parallels with processes described in amphibians, where retinal regeneration can arise from non-neural sources such as the retinal pigmented epithelium (RPE). For example, in newts or *Xenopus*, differentiated RPE cells undergo dedifferentiation and epithelial-to-mesenchymal transition, acquire progenitor-like features, and subsequently transdifferentiate into multiple retinal neuronal classes without forming a persistent glial scar^70–74^. While this process is limited or absent in other vertebrates such as mammals, it is conserved in chicken during embryogenesis^75–78^. Together, these observations support the existence of evolutionarily conserved regenerative modules in vertebrate eyes in which neural-crest- and/or mesenchymal-derived populations can be mobilized to regenerate retinal neurons through transdifferentiation rather than classical stem-cell-driven neurogenesis.

Together, these findings have important translational implications. In mammalian retinae, Müller glia exhibit limited regenerative potential and are constrained by epigenetic and inflammatory barriers. Our identification of a mesenchymal-derived regenerative pathway suggests that alternative endogenous cell populations, particularly neural crest-derived fibroblast-like cells, may represent underexplored reservoirs of plasticity supporting tissue repair. Targeting Pdgf, Wnt, Tgf-β, Hippo, and/or insulin signaling could provide a framework for inducing controlled mesenchymal reprogramming *in situ* to support tissue and organ regeneration, potentially circumventing some of the limitations associated with stem-cell transplantation or *in vivo* glial reprogramming.

In summary, this study reveals a previously unrecognized regenerative axis in the zebrafish retina in which mesenchymal-derived neural progenitors participate to the process of neuronal regeneration. These results broaden the cellular landscape of vertebrate retinal regeneration and highlight mesenchymal plasticity as a promising avenue for developing endogenous neural-repair strategies in the mammalian central nervous system, with potentially broader implications for tissue regeneration in other organs. Future studies will aim to define whether such strategies can be translated into cell therapy for the treatment of glaucoma and other retinal degenerative diseases in human.

## MATERIALS AND METHODS

### Study design and statistical analysis

Research subjects (larval stage and genotype) and treatments applied are defined in the figures and/or figure legends. Experimental analyses were performed with a minimum of 3 biological replicates (and a minimum of 3 technical replicates when applicable). Regarding statistical power, we calculated the effect size with Cohen’s D and found that all measured differences had D-values that exceeded an absolute value of 4, which qualifies as a very large effect. For experiments using zebrafish larval retina as one biological replicate, a minimum of 3 larvae (6 retinae) were used. The number of biological replicates and larvae used for each specific experiment as well as the statistical test used to calculate significance is indicated in the figure legends. For statistical analysis, p-values are indicated in the figures. Comparisons were conducted to determine significant differences between control and experimental conditions. Statistical analyses were performed using GraphPad Prism 10. The normality of data distribution was assessed using the Shapiro-Wilk test and visualized with quantile-quantile (Q-Q) plots. Homogeneity of variance between groups was evaluated using F-tests. Based on these preliminary analyses, Student’s T-test, T-test with Welch’s correction, one-sample Wilcoxon test, Mann-Whitney test, or Fisher’s exact test were selected for subsequent statistical comparisons, as indicated in the figure legends. Statistical significance was set at p < 0.05.

### Zebrafish lines and developmental conditions

Embryos were raised and staged according to standard protocol^79^. Experiments were performed in accordance with animal-care guidelines. Our IACUC animal protocol (AUP#20-06) is approved by the MDI Biological Laboratory. Transgenic zebrafish lines used in this study are: Tg(*islet2b*:gal4)^36^; Tg(*uas*:NTR2.0-YFP)^35^; Tg(*uas*:NTR2.0-mCherry)^35^; Tg(*ins*:NTR2.0-mCherry)^80^; Tg(*mpeg*:mCherry)^81^; Tg(*pdgfrb*:mCherry)^82^; Tg(*pdgfrb*:H2B-dendra2)^83^; Tg(*sox10*:Kaede)^84^; Tg(*sox10*:creERT2)^85^; Tg(*hsp70*:Lox-DsRed-Lox-GFP)^86^; and Tg(*gfap:NTR-tagRFP*). Mutant zebrafish lines used are the *pdgfrb*^87^ and *sox10* mutants, the latter of which we generated in this study (see below). Photoconversion was performed using UV light in the incubator for whole-larvae photoconversion. For imaging analysis, larvae were fixed overnight at 4°C in 4% paraformaldehyde/1xPBS, after which they were dehydrated through an ethanol series and stored at −20°C until use.

### Plasmid construction and transgenic-line establishment

To establish stable transgenic lines, plasmids were injected into one-cell-stage embryos with Tol2 transposase mRNA^88^. Generation of the Tg(*gfap:NTR-tagRFP*) line was performed using 10 kb of the *gfap* promoter cloned in the p5E entry vector. The NTR-tagRFP coding sequence was cloned in the middle-entry vector. The appropriate entry and middle-entry clones were mixed with the SV40pA 3’ entry vector and recombined into the Tol2 transposon destination vector.

### CRISPR/Cas-9 generation of a zebrafish *sox10* mutant

To generate a *sox10* mutant, we took advantage of a previously described genome-editing method^89^. Specifically, we used tracrRNA and crRNA (IDT) to form functional gRNA duplexes targeting the ORF (5’-CACCCCCAAGACGGAACTGC-3’ and 5’-GAAGATGACCGGTTCCCCAT-3’). Co-injection of these gRNAs with the Cas-9 protein (IDT) resulted in indels in the coding sequence, identified by Sanger sequencing and analyzed with ICE software (Synthego) in F0 injected embryos. F0 carriers for mutations in the *sox10* locus in the germline were identified by sequencing the alleles on the clutches, and were then outcrossed with Tg(*islet2b*:gal4); Tg(uas:NTR2.0-YFP) lines to obtain F1 heterozygous mutants. We identified F2 homozygous mutants via the presence of a *sox10* loss-of-function phenotype^90^.

### Pharmacological treatments

Stock solution of 1-phenyl 2-thiourea (PTU; Sigma) was prepared at 25X (0.075%). PTU 1X working solution was used to inhibit pigmentation in zebrafish embryos. A stock solution of hydroxytamoxifen (4-OHT; Sigma) was prepared at 10 mM in 100% DMSO (Sigma). For induction of LoxP-site recombination, zebrafish larvae were treated with 4-OHT at a concentration of 10 µM, after which we performed heat-shock for 1 hour at 37.5°C to induce fluorophore expression. The larvae were then left in the incubator for 1 hour at 28.5°C for recovery before fixation. Genetic cell ablation was performed using metronidazole (MTZ; Sigma) at a concentration of 10 mM. Inhibition of cellular processes was performed using specific Pdgf (Sigma; PDGFR Tyrosine Kinase Inhibitor V, 521234; 2 μM), Wnt (Sigma; IWR1, I0161; 10 μM), Tgf-β (Tocris; SB431542; 10 μM), Bmp (Tocris; DMH-1; 10 μM), Hippo (Sigma; Verteporfin, SML0534; 10 μM), and insulin (Sigma; HNMPA-(AM)3, 397100; 20 μM) signaling inhibitors or using blebbistatin (Tocris; 10 μM) for Myosin II inhibition. CHIR99021 (Sigma; 20 μM) was used as a potent and selective activator of Wnt signaling pathway (

### Gene-expression analysis

*In situ* hybridizations were performed as previously described^91^. The ORFs of *sox10, pdgfra, pdgfrb, col2a1, col12a1*, and *insulin* were cloned in a pCS2+ vector using zebrafish cDNA. Antisense DIG-labelled probes were transcribed using T3 polymerase (Promega) and the linearized pCS2+ plasmid containing the ORF as a template. DIG probes were detected using anti-DIG antibody (Roche), and *in situs* were revealed using BCIP reagent (Roche). HCR fluorescent *in situ* experiments were performed using probes, fluorophores, and protocols provided by Molecular Instruments (HCR™ RNA-FISH (v3.0)).

Single-cell RNA sequencing datasets used in this study are from Emmerich et al., 2024, and available under accession number GSE268179^36^. Enrichment with a known specific zebrafish marker gene was used to identify cell populations in the subset analysis: neural progenitors (*ascl1*a, *ascl1b*, *neurog1*, *neurod1*, or *neurod4*), RGC precursor cells (*atoh7*, *islet2b*, *pou4f1*, *pou4f2*, or *pou4f3*), post-mitotic neurons (*elavl3* or *elavl4*), neural/glial cells (*olig2*), myelinated glial cells (*mbpa* or *mbpb*), neural crest/glial cells (*sox10*), and vascular cells (*kdrl* or *fli1a*). Using SeuratV4^92^, UMAP representations were generated for subsequent pseudo-time and velocity analysis. For trajectory analysis, each group cluster was isolated and pseudo-time plotting of transition was generated using the Monocle package^93^. As a metric of cell activity, we applied the VelocityR package that assesses the relative gene velocity, indicative of cell activity and change in direction^94^. Differential expression analysis was performed to identify genes differentially expressed in *sox10*+ retinal cells of interest, compared against all other cells in the sub-cluster.

Total RNA samples for bulk RNAseq analysis were prepared with ∼50 retinae/sample. Each experimental condition was performed using biological replicates to ensure reproducibility. Trizol (Invitrogen) was used to homogenize cellular extracts with the TissueLyzer (Qiagen). RNA purification was performed as described above. For bulk RNAseq analysis, library preparation and sequencing were carried out by NovoGene using the Illumina NGS platform. Indexing and alignment were performed using HISAT2 (v2.0.5)^95^ against the GRCz11 (*Danio rerio*) reference genome obtained from Ensembl. Mapped reads were assembled using StringTie (v1.3.3b)^96^, and quantification was performed using FeatureCounts (v1.5.0-p3)^97^. Differential expression analysis was conducted in the R environment using DeSeq2^98^, and Gene Set Enrichment Analysis (GSEA) was performed using clusterProfiler^99^. All visualizations were generated using ggplot2.

### Immunostaining

Immunostaining was performed using anti-GFP (1/1,000, Torrey Pines Biolabs), anti-DsRed (1/500, Takara), anti-HuC/D (1/500, Invitrogen), anti-acetylated tubulin (1/500, Sigma) anti-Insulin (1/250, GeneTex) as primary antibodies and Alexa 405, Alexa 488, Alexa 555, or Alexa 647-conjugated goat anti-rabbit immunoglobulin G (IgG) or goat anti-mouse IgG (1/1,000) as secondary antibodies (Invitrogen). Click-iT EdU Cell Proliferation Assays kit was used to assess cell proliferation (Invitrogen).

### Image acquisition

Images were acquired using a point scanning confocal unit (LSM 980, Carl Zeiss Microscopy, Germany) on a Zeiss Axio Examiner Z1 upright microscope stand (409000-9752-000, Carl Zeiss Microscopy, Germany). The images underwent modifications using Fiji software, and quantification was performed using Fiji or Imaris software.

## Supporting information

SuppFigures

SuppTableS1

## ACKNOWLEDGMENTS

This study was supported by Institutional Development Awards (IDeA) from the National Institute of General Medical Sciences of the National Institutes of Health under grant numbers P20GM103423 (MDIBL), P20GM104318 (MDIBL), and P20GM144265 (RM). Funding was also provided by the Morris Discovery Fund (RM). This work was also funded by the following grants from the National Institutes of Health (F31 EY032790-01 to K.E., P30EY001765-45 to the Wilmer Eye Institute, T32EY007143-24 to K.E., and R01EY026580 to J.S.M), and by a Brightfocus Foundation National Glaucoma Research grant (G2020315 to J.S.M.).

Image collection, processing, and analysis for this manuscript was performed with the assistance of Dr. Frederic Bonnet and the MDI Biological Laboratory’s Light Microscopy Facility (RRID:SCR_019166), which is supported by an Institutional Development Award (IDeA) from the National Institute of General Medical Sciences of the National Institutes of Health under grant number P20GM103423.

## Authors’ contributions

*Conceptualization*: Bidhi Diwedi, Anindita Neog, Aissette Baanannou, Jeff S. Mumm, and Romain Madelaine. *Investigation*: Bidhi Diwedi, Anindita Neog, Aissette Baanannou, Pritha Das, Caroline Halluin, James H. Thierer, Michael Patnaude, and Romain Madelaine. *Data curation*: Bidhi Diwedi, Anindita Neog, Aissette Baanannou, Pritha Das, Romain Menard, Dexter Morse, Kevin Emmerich, Joel Graber, and Romain Madelaine. *Formal analysis*: Bidhi Diwedi, Anindita Neog, Aissette Baanannou, Pritha Das, Romain Menard, Dexter Morse, Frederic Bonnet, Joel Graber, and Romain Madelaine. *Funding acquisition*: Jeff S. Mumm and Romain Madelaine. *Supervision*: Romain Madelaine. *Writing – original draft:* Romain Madelaine.

We thank Drs. N.D. Lawson, E.C. Liao, A. Siekmann, and D. Whener for kindly sharing transgenic and mutant lines.

## Data availability

All data, reagents, and genetic tools presented in this manuscript are available to the scientific community.

## FIGURE LEGENDS

**Figure S1. RGC genetic ablation induces a regenerative response in the zebrafish retina. (A)** Sub-analysis of single-cell RNAseq control and ablated datasets at 6 dpf after genetic ablation of RGCs using Tg(*islet2b:gal4*); Tg(*uas:NTR-yfp*) with MTZ treatment between 5 and 6 dpf. UMAPs show the clustering of various retinal cell types. Each cluster is color-coded to represent different cells as indicated in the figure. These panels underscore the fate proximity of *sox10*+ cells (green) with neural progenitor cells (blue) in two clusters (red arrows). **(B)** Trajectory (left panel) and cell velocity (right panel) in cluster 2 reveal a pattern of movement from *sox10*-expressing cells to young and mature neurons. **(C)** Differential gene expression analysis of the *sox10*+ cells against all other cells in the cluster 2, indicating genes significantly associated with *sox10* expression in cluster 2. **(D-F)** Genetic ablation of RGCs using Tg(*islet2b:gal4*); Tg(*uas:NTR-yfp*) with MTZ treatment between 5 and 6 dpf leads to optic-nerve degeneration (D), RGC death as revealed by reduced acetylated tubulin staining (E), and increased microglia presence in the injured GCL (F). **(G)** Representative image (confocal z-section) of Tg(*pdgfrb:mCherry*) and HuC/D expression in the GCL after RGC genetic ablation (between 5 and 6 dpf). **(H)** Representative image (confocal z-projection) of Tg(*sox10:kaede*) expression and EdU incorporation in the GCL after RGC genetic ablation (between 5 and 6 dpf). Scale bars: 100 μm.

**Figure S2. *sox10*/*pdgfrb*-expressing cells also carry a fibroblast identity. (A)** Representative image (confocal z-section) of *pdgfrb* and *olig2* co-expression in the GCL after RGCs genetic ablation (between 5 and 6 dpf) using HCR fluorescent *in situ hybridization*. **(B)** Representative image (confocal z-section) of *pdgfrb* and *neurod1* co-expression in the GCL after RGC genetic ablation (between 5 and 6 dpf) using HCR fluorescent *in situ hybridization*. **(B’)** Quantification of the number of *pdgfrb*+ cells in the GCL at 6 dpf after genetic ablation of RCC between 5 and 6 dpf (control; n=5 and MTZ; n=9; t-test with Welch’s correction, left). Analysis of the percentage of *pdgfrb*+ cells expressing the proneural factors *neurod1* (control; n=5 and MTZ; n=9; one-sample Wilcoxon test; right). **(C)** Representative image (confocal z-projection) of sox10:kaede (green) *and* PC_sox10:kaede (purple) without injury showing the transition from green to red fluorescence of the Kaede protein in the eye after UV photoconversion of larvae. **(D)** Representative image (confocal z-section) of *sox10* or *pdgfrb* expression in the GCL after RGC genetic ablation (between 5 and 6 dpf) using HCR fluorescent *in situ hybridization* in the *pdgfrb* (top) or *sox10* (bottom) mutant, respectively. **(D’)** Quantification of the number of *sox10* or *pdgfrb*+ cells in the GCL at 6 dpf after genetic ablation of RCCs between 5 and 6 dpf in the *pdgfrb* (control; n=12; MTZ; n=13; and MTZ in *pdgfrb* mutant; n=12; Fisher’s exact test; left) or *sox10* (control; n=21; MTZ; n=21; and MTZ in *sox10* mutant; n=21; Fisher’s exact test; right) mutant, respectively. **(E)** Representative image (confocal z-section) of *pdgfrb* and *acta2* co-expression in the GCL after RGC genetic ablation (between 5 and 6 dpf) using HCR fluorescent *in situ hybridization*. Error bars represent s.d. Scale bars: 100 μm.

**Figure S3. Signaling pathways involved in fibroblast activation after injury participate in the accumulation of *sox10*-derived progenitors in the GCL. (A)** Analysis of *sox10* mRNA expression by *in situ* hybridization in control or MTZ-treated (between 5 and 6 dpf) larvae (Tg(*islet2b:gal4*); Tg(*uas:NTR-mCherry*)) and/or Tg(*gfap:NTR-tagRFP*) at 6 dpf. Quantification of the number of retina harboring *sox10*+ cells in the GCL at 6 dpf after genetic ablation of RGCs and/or *gfap*+ cells between 5 and 6 dpf using Fisher’s exact test (control; n=10 MTZ (islet2b); n=11; MTZ (gfap); n=13; MTZ (gfap + islet2b; n=10). **(B)** Representative image (confocal z-section) of *pdgfrb* and *eys2* co-expression in the GCL after RGC genetic ablation (between 5 and 6 dpf) using HCR fluorescent *in situ hybridization*. **(C)** Analysis of *sox10* mRNA expression by *in situ* hybridization in control or MTZ-treated (between 5 and 6 dpf) larvae (Tg(*islet2b:gal4*); Tg(*uas:NTR-mCherry*)) at 6 dpf with or without Tgf-β or Bmp signaling inhibition, or with or without myosin II activity inhibition with blebbistatin. **(C’)** Quantification of the number of retina harboring *sox10*+ cells in the GCL at 6 dpf after genetic ablation of RCCs between 5 and 6 dpf using Fisher’s exact test (control; n=15; MTZ; n=26; MTZ + Tgf-β inhibitor; n=28; MTZ + Bmp inhibitor; n=47; MTZ + blebbistatin; n=29). **(D)** Representative image (confocal z-section) of *pdgfrb* expression in the GCL after RGC genetic ablation (between 5 and 6 dpf) with or without the Wnt signaling pathway activator (CHIR99021) using HCR fluorescent *in situ hybridization*. Arrows show *pdgfrb*+ cells outside of the GCL. Scale bars: 100 μm.

**Figure S4. Identification of *insulin* as locally expressed in the retina after ablation of RGCs. (A)** HCR analysis revealed co-expression of the *pgdfrb* and *insulin* genes in the GCL after ablation of RGCs (confocal z-section). **(B, B’)** GO (Gene Ontology) and KEGG analysis between control and MTZ-treated larvae at 6 dpf. **(C)** Analysis of *insulin and sox10* mRNA expression by *in situ* hybridization in control- or MTZ-treated (between 5 and 6 dpf) larvae (Tg(*islet2b:gal4*); Tg(*uas:NTR-mCherry*)) and/or Tg(*ins:NTR-mCherry*) at 6 dpf. **(C’)** Quantification of the number of retina harboring *sox10*+ cells in the GCL at 6 dpf after genetic ablation of RGCs and/or *insulin*+ cells between 5 and 6 dpf using Fisher’s exact test (control; n=10; MTZ (islet2b); n=10; MTZ (insulin + islet2b; n=10). **(D)** Representative image (confocal z-section) of *pdgfrb* and *insr* gene expression in the retina in the control condition and after RGC genetic ablation (between 5 and 6 dpf) using HCR fluorescent *in situ* hybridization. Scale bars: 100 μm.

**Table S1. List of dysregulated genes at 6 dpf after RGC ablation between 5 and 6 dpf.**

