## Supplementary figures and images for "Mesenchymal-derived neural progenitors underlie local *insulin* production and neuronal transdifferentiation during retina regeneration"

### SuppFigures

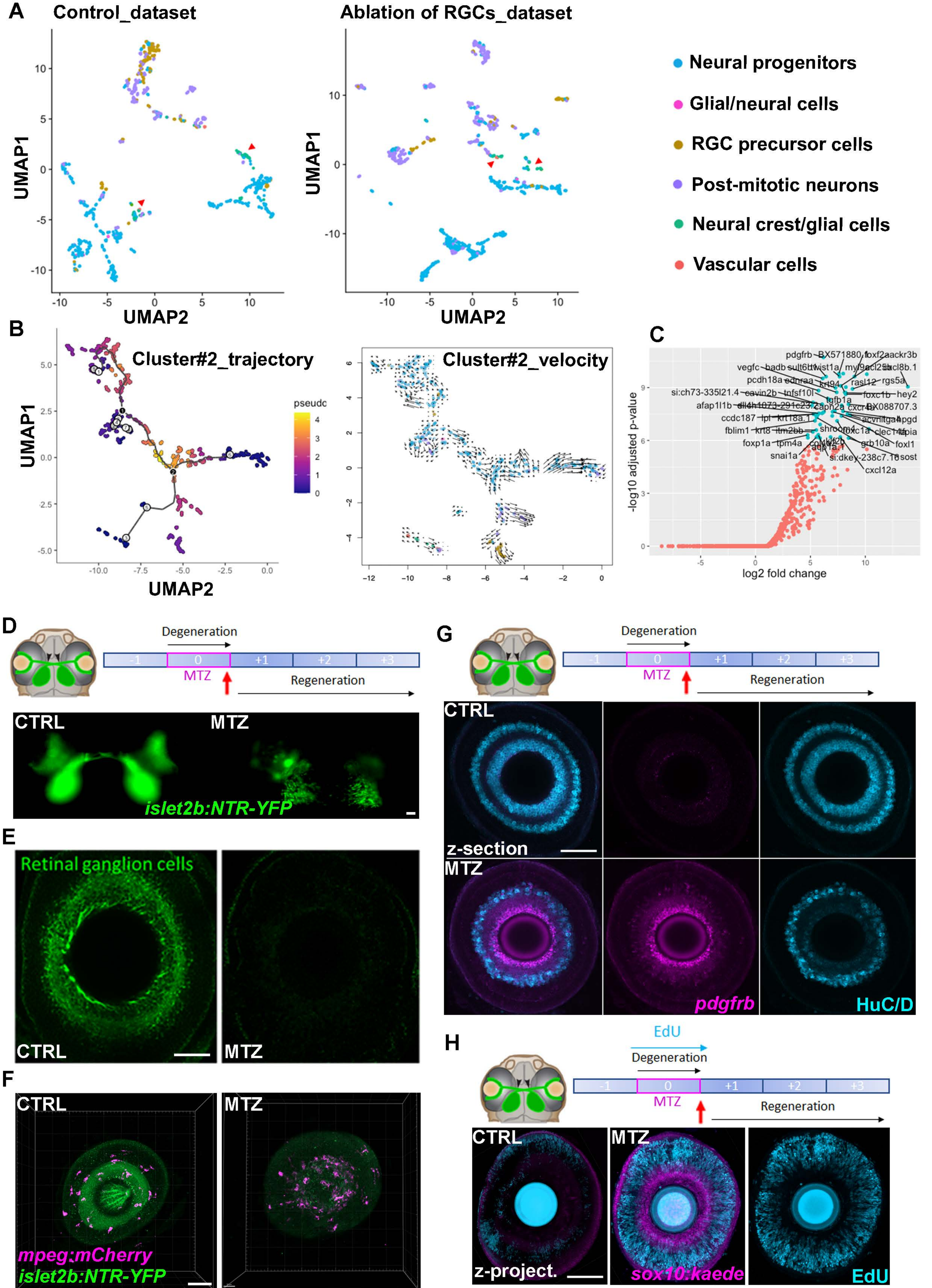

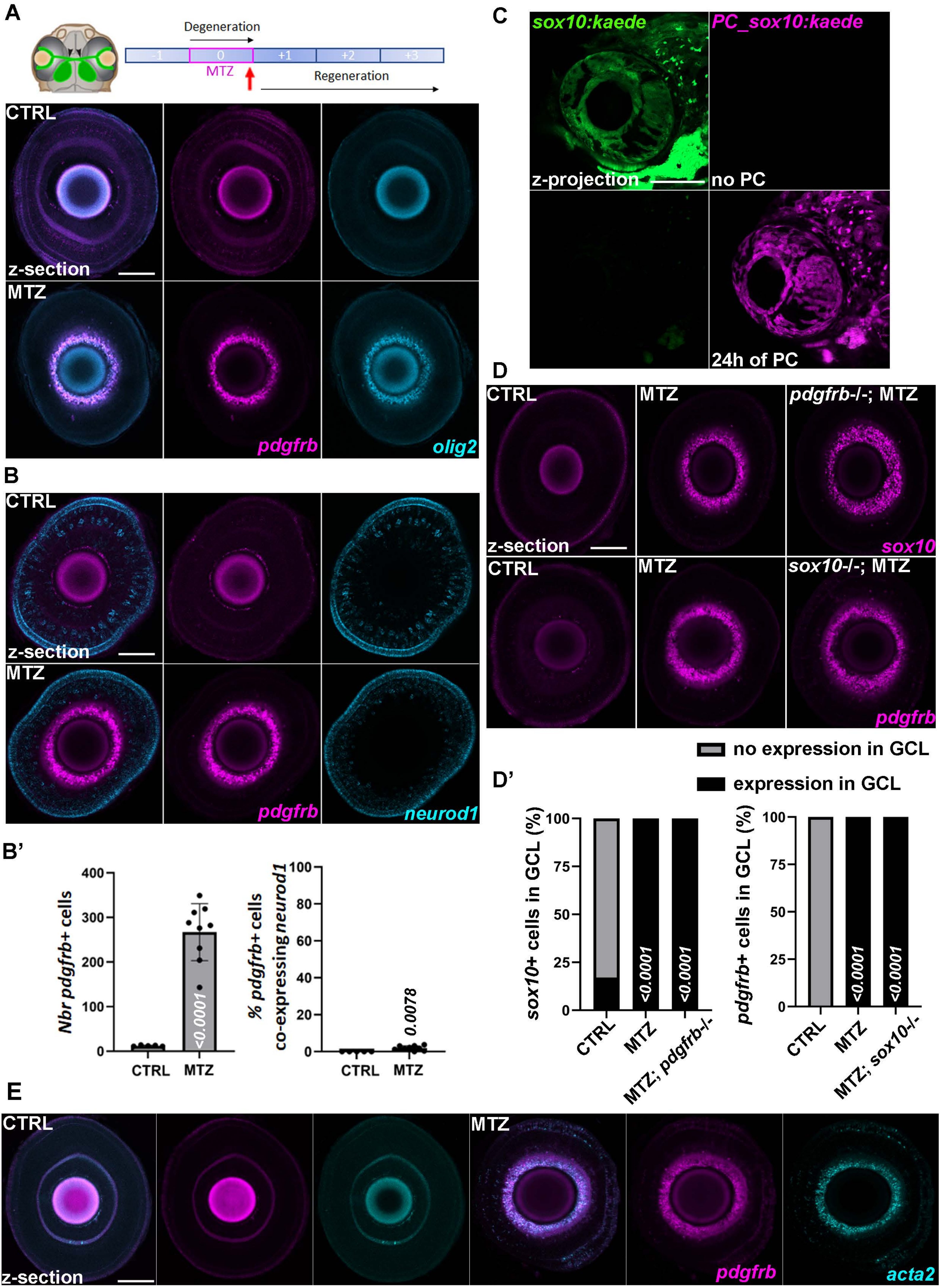

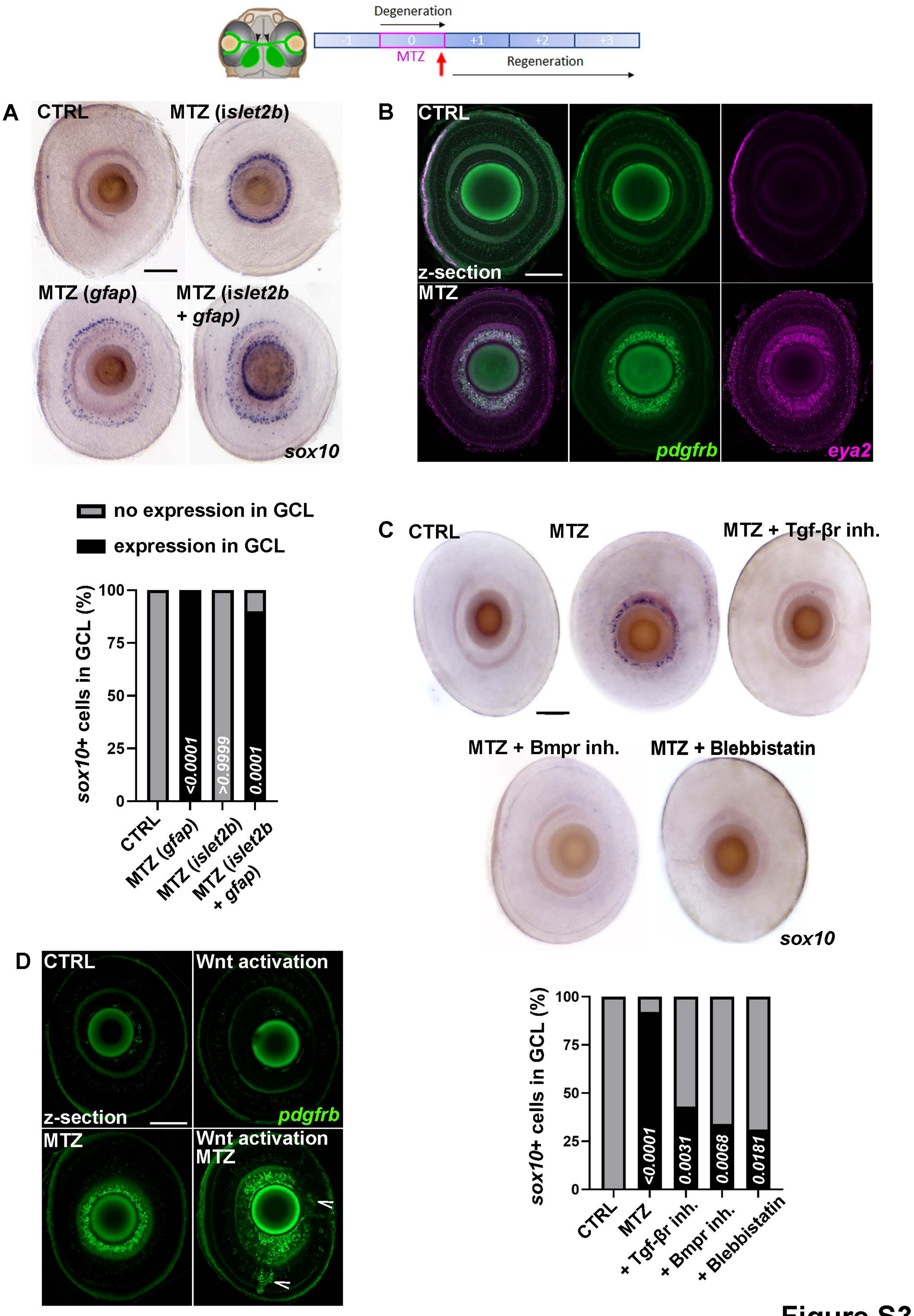

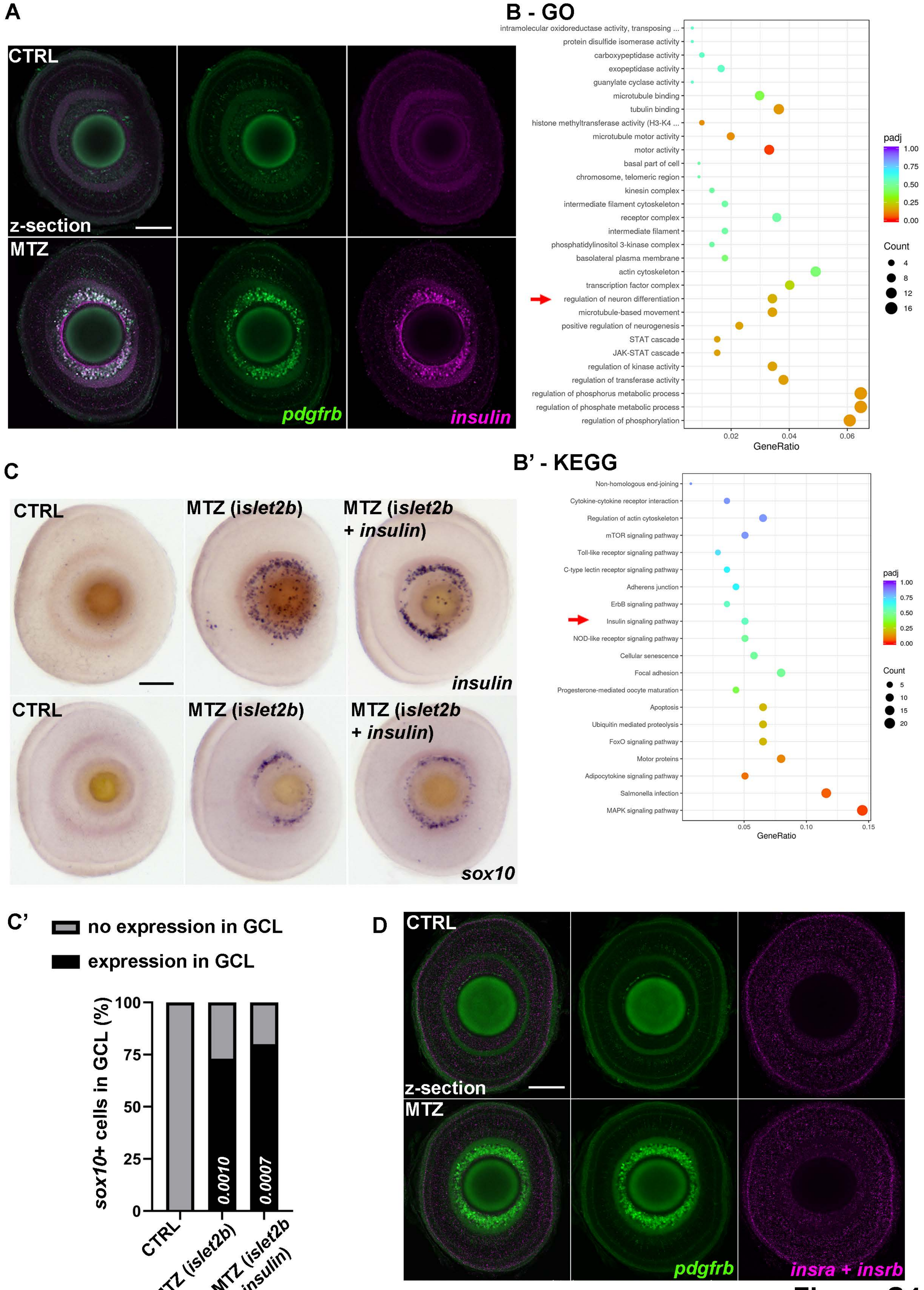
